# Integrating multiple dimensions of biodiversity to inform global parrot conservation

**DOI:** 10.1101/812321

**Authors:** Kevin R. Burgio, Katie E. Davis, Lindsay M. Dreiss, Laura M. Cisneros, Brian T. Klingbeil, Steven J. Presley, Michael R. Willig

## Abstract

Because biodiversity is increasingly threatened by habitat destruction and climate change, conservation agencies face challenges associated with an uncertain future. In addition to changes associated with climate and land use, parrots are threatened by hunting and capture for the pet trade, making them the most at-risk order of birds in the world. Parrots provide key ecosystem services, but remain understudied compared to other major bird orders despite their high extinction risk and ecological importance. Species richness is often used to identify high priority areas for conserving biodiversity. By definition, richness considers all species to be equally different. However, ongoing research emphasizes the importance of incorporating ecological functions (functional diversity) or evolutionary relationships (phylogenetic diversity) to more fully understand patterns of biodiversity, suggesting that using functional and phylogenetic information could improve conservation strategies. These distinctions among dimensions of biodiversity are important, because (1) areas of high species richness do not always represent areas of high functional or phylogenetic diversity, and (2) functional or phylogenetic diversity may better predict ecosystem function and evolutionary potential, which are essential for effective long-term conservation policy and management.

Our objective was to create a framework for identifying areas of high species richness, functional diversity, and phylogenetic diversity within the global distribution of parrots We combined species richness, functional diversity, and phylogenetic diversity into an Integrated Biodiversity Index (IBI) to identify global biodiversity hotspots for parrots. We found important spatial mismatches among dimensions, which demonstrate that species richness is not always an effective proxy for other dimensions of parrot biodiversity. The IBI is an integrative and flexible index that can incorporate multiple dimensions of biodiversity, resulting in an intuitive and more direct way of assessing comprehensive goals in conservation planning (i.e., healthy ecosystem functioning and climate change resilience).

## INTRODUCTION

As we enter the early stages of the “Sixth Mass Extinction” (Ceballos et al. 2015), conservation agencies are struggling to face the challenges of a less certain future (Armsworth et al. 2015) as a consequence of habitat conversion and climate change (Urban et al. 2015). While most taxa are at risk, parrots (Psittaciformes) are especially so, as they are the most threatened order of birds, with ~43% of species listed as near-threatened or worse, and with ~28% listed as threatened or worse by the IUCN (IUCN 2015, Olah et al. 2016). Furthermore, 56% of parrot populations are currently thought to be in decline (IUCN 2015). However, estimates of bird density exist for only 25% of species, and those estimates suggest that parrot density is higher inside of protected areas than outside of them (Marsden & Royle 2014), demonstrating the critical importance of conservation action.

Parrots provide many important ecosystem functions, including invertebrate pest management, pollination, seed dispersal, and genetic-linking of plant communities, making them “keystone mutualists” (Tella et al. 2015, Blanco et al. 2015, Blanco et al. 2016). Parrots also forage on plants that are toxic and poisonous to many vertebrate species (Gilardi & Toft 2012, Blanco et al. 2015), allowing them to find food and persist in habitats where other frugivorous species cannot (Gilardi & Toft 2012). Given the critical ecosystem services parrots provide, their loss may have detrimental effects on the persistence of many plant species, thereby contributing to ecosystem instability. Conversely, some parrot species have been spreading into new areas, primarily as a consequence of the pet trade, which has led to human/wildlife conflicts in Europe (White et al. 2019) and the introduction of 25 non-native species that now breed in the USA (Uehling et al. 2019).

In general, parrots have long generation times, low population densities, and high risk of being hunted or trapped (Pires 2012, Marsden & Royle 2014), all characteristics associated with high extinction risk (Bennett and Owens 1997, Cardillo et al. 2005). Predictably, parrot species that are larger-bodied and have longer generation times are generally found at relatively lower densities (Marsden & Royle 2014) and are more likely than are smaller parrot species to be obligate tree-cavity nesters (Renton et al. 2015), increasing extinction risk (Jones et al. 2006, Olah et al. 2016). Despite being the most-threatened order of birds, parrots are relatively understudied compared to other orders of birds (Brito & Orpea 2009, Ducatez & Lefebvre 2014). The lack of research may explain why density estimates, one of the most important factors in determining conservation status (Mace et al. 2008), are inadequate for most parrot species (Marsden & Royle 2014).

In addition to the effects of climate change and habitat loss resulting from agriculture or logging, other anthropogenic activities such as hunting and trapping are among the biggest threats to parrots (Snyder et al. 2000, Olah et al. 2016). Logging and agricultural conversion of habitats disproportionately affect parrots because a majority of parrot species (~70%) are forest-dependent (Olah et al. 2016). Most parrot species (~78%) also rely on tree-cavities for nesting (Renton et al. 2015), and nesting in some species is limited to a few tree species (Renton and Brightsmith 2009).

Over the past few decades, one of the most common methods for identifying areas of highest conservation priority has been based on the “hotspot concept” (Reid 1997, Myers et al. 2000), which uses existing species range maps to prioritize conservation efforts where species richness or richness of endemic species is highest. More recently, the multidimensional nature of biodiversity has emerged as a critical consideration for conservation (e.g., Isaac et al. 2007, Mazel et al. 2014, Brum et al. 2017, Pollock et al. 2017) and considerations of irreplaceability and complementarity have supplemented the hotspot concept via spatial prioritization approaches (e.g. Andelman and Willig 2002, 2003; Zonation and Marxan; reviewed in Moilanen et al. 2009). Regions of high functional diversity (a measure of ecological trait diversity within an assemblage) or phylogenetic diversity (a measure of evolutionary diversity within an assemblage) may not coincide with areas with high species richness (e.g. for mammals; see Safi et al. 2011, Mazel et al. 2014, Brum et al. 2017). Consequently, calls to incorporate phylogenetic or functional biodiversity into conservation planning have arisen over the last couple of decades (Mace et al. 2003, Diaz et al. 2007).

Limited availability of data at appropriate scales, and the complex nature of metrics used to quantify functional or phylogenetic diversity, have inhibited until very recently the incorporation of this information into conservation planning approaches that prioritize land for protection (e.g. Brum et al. 2017, Pollock et al. 2017). Phylogenetic or functional diversity may be better indicators of community resilience than is species richness (Naeem et al. 2012). Conserving functional diversity is critical for maintaining ecosystem functions (Naeem et al. 2012), and thus for maintaining critical ecosystem services, making it an important consideration in conservation (Chan et al. 2006, Diaz et al. 2007, Cimon-Morin et al. 2013, Kosman et al. 2019). Nonetheless, cases that incorporate functional diversity in conservation research are rare (but see Devictor et al. 2010 and Mazel et al. 2014). Maintaining the capacity for future adaptation is an important consideration for communities undergoing rapid climatic and environmental changes, suggesting that phylogenetic diversity should be given consideration when determining conservation goals (Naeem and Li 1997, Cardinale et al. 2012). Moreover, the loss of phylogenetic or functional diversity may be a better indicator than is the loss of species richness in quantifying ecosystem vulnerability (Srivastava et al. 2012).

Here, we created a framework to identify areas of high species richness, functional diversity, and phylogenetic diversity within the global distribution of parrots. We separately calculated species richness, functional diversity, and phylogenetic diversity, and then combined them into an Integrated Biodiversity Index (IBI) to identify global biodiversity hotspots to aid in parrot conservation. Important spatial mismatches between dimensions indicate situations in which species richness is not an effective proxy for other dimensions of parrot biodiversity.

## METHODS

### Data collection

#### Distribution data

We used range maps for all 398 extant species of parrot (Birdlife International 2015) following the taxonomy of del Hoyo et al. (2014) to inform spatially-explicit estimates of biodiversity at a global scale.

#### Trait and phylogenetic data

We estimated functional diversity using two types of data: categorical (binary) and mensural traits (Table 1). For each data type, we used a suite of traits that reflect particular niche axes and define functional components. Categorical traits included components of diet, foraging strategy, and foraging location, whereas mensural traits comprise body size and range size. For each categorical trait, a species received a “1” if it exhibited the characteristic and a “0” if it did not. For each body size, we used the average value for each species based on measurements of multiple adults, when available. We obtained trait data for all parrot species from the literature (see Burgio et al. [2019] for more details) and range size data from Birdlife International (2015). We calculated phylogenetic diversity for each community using branch lengths found in a recently-published time-calibrated phylogenetic supertree for all 398 extant parrots (Burgio et al. 2019).

**Table 1:**
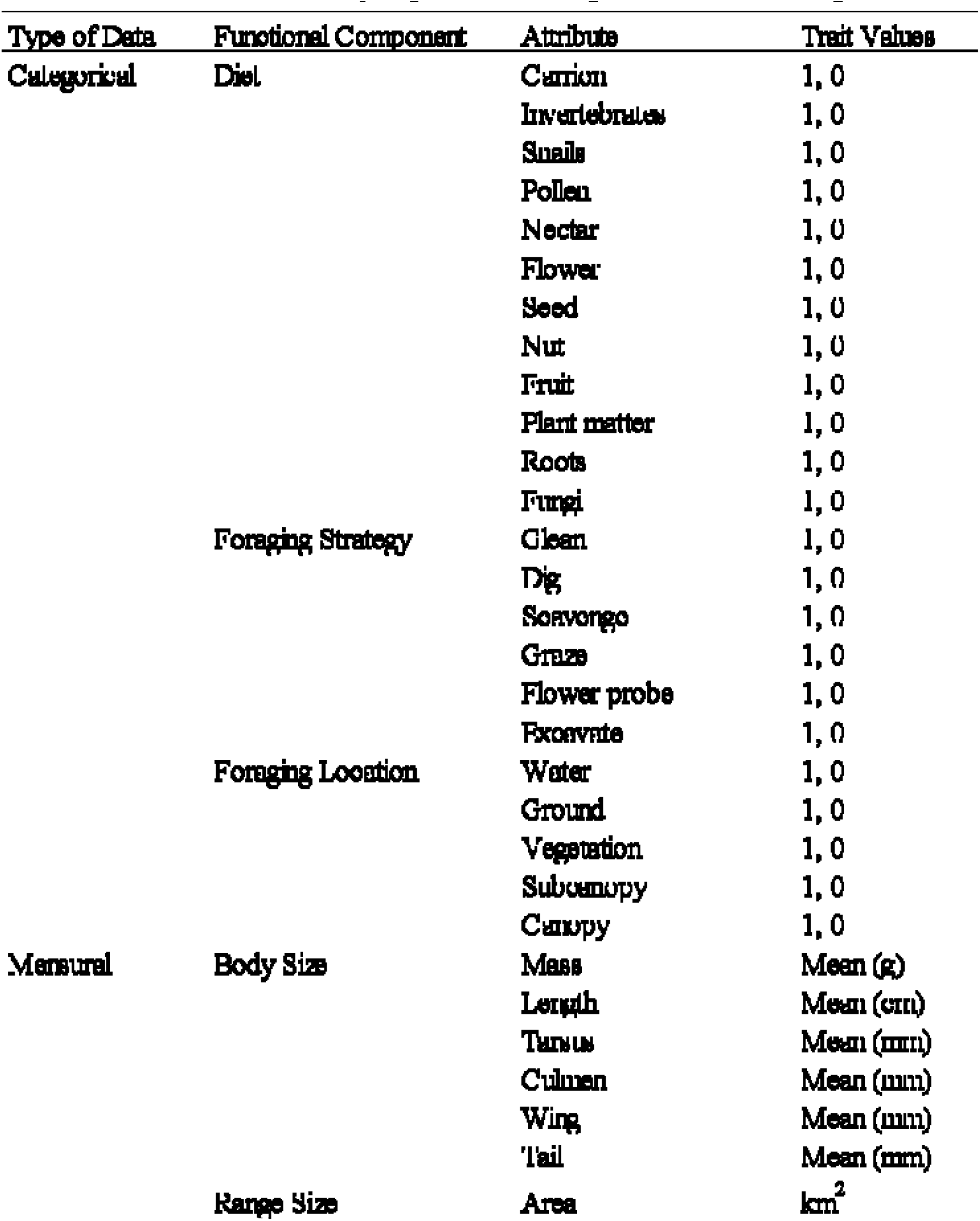
Functional attributes that reflect micho axes (functional components) were used to estimate functional biodiversity of parrot assemblages for each 500 km^2^ grid coil.

### Analyses

#### Biodiversity indices

We created a grid in ArcMap v.10.3 (ESRI, Redlands, CA, USA), using the Cylindrical Equal Area projection, with each grid cell measuring 50 x 50 km (hereafter “grid cell”). For each grid cell (*n* = 21,078), we estimated species richness as the number of species with a range overlapping the cell. We estimated phylogenetic and functional diversity for each cell using Rao’s quadratic entropy (Rao’s Q; Botta-Dukát 2005). Rao’s Q measures the average difference between all pairs of species, thereby reflecting multivariate dispersion. We obtained the average phylogenetic or functional distances among species from pairwise dissimilarity matrices for the phylogenetic and functional components, as well as separately for each of the six functional categories. For the phylogenetic supertree, we populated a pairwise dissimilarity matrix via the “cophenetic” function of the R package “ape” (v.3.5, Paradis et al. 2004). We used the Gower metric from the R package “cluster” (v.2.0.4, Maechler et al. 2012) to calculate pairwise functional dissimilarity matrices.

To allow meaningful comparisons among dimensions, we transformed each metric into its effective number of species or Hill number (hereafter numbers equivalent). The numbers equivalent is the number of maximally dissimilar species that is required to produce an empirical value of a diversity metric (Jost 2006). This transformation facilitates intuitive interpretation of differences among assemblages and dimensions because indices are expressed in the same units (Jost 2006, Chao et al. 2014). Species richness is its own numbers equivalent. We transformed Rao’s Q values into numbers equivalents using R functions developed by de Bello et al. (2010).

#### Integrated Biodiversity Index

The Integrated Biodiversity Index (IBI) combines numbers equivalent transformations of Rao’s Q for phylogenetic and functional diversity (all traits combined) with species richness. We scaled each dimension of biodiversity to a range from 0 to 1 so that each would have equal opportunity to contribute to the IBI value. Without such scaling, species richness would likely dominate spatial patterns of biodiversity. The IBI is the sum of the scaled representations of species richness (S), functional diversity (FD), and phylogenetic diversity (PD) for a particular grid cell (*i*):

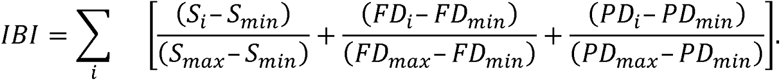

As a consequence of the numbers equivalent transformation and scaling functions, IBI values range from 0 to 3 and equally weight each dimension of diversity (i.e. a value of “0” would mean low combined biodiversity while a value of “3” would be highest in combined biodiversity).

## RESULTS

Species richness of parrots is highest in the Amazon Basin of South America, along the southeastern coast of Australia, and in the mountainous region of New Guinea (Fig. 1a). Functional diversity is highest in the dry Chaco of South America (Fig. 1b). Our measure of functional diversity represents multivariate dispersion, which is greatest for assemblages that represent many functional types, but that have low redundancy in those functions. Dry Chaco parrot assemblages have low species richness (Fig. 1a) and species that differ greatly from each other in functional traits associated with diet and foraging location. Phylogenetic diversity is highest in Australia, arising primarily from the diversification of multiple subfamilies within the Psittacidae, and the fact that cockatoos (Cacatuidae), which represent a deep split in the parrot phylogeny (Fig. 2), are endemic to Australia and Oceania.

**Figure 1.**
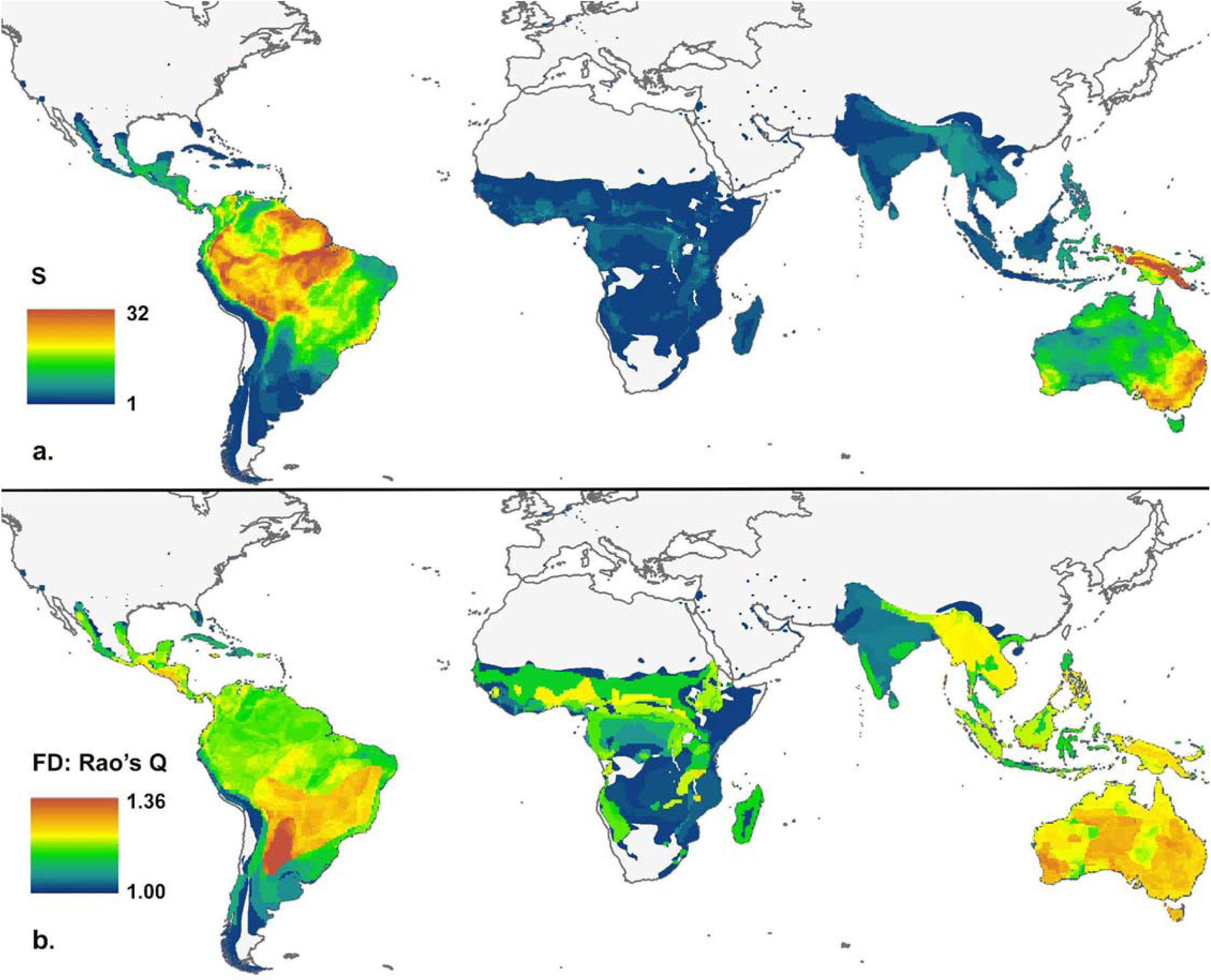
Global map of parrot (a) species richness (S) and (b) functional diversity (FD; Rao’s Q, based on Hill numbers). Functional traits in the analysis include: diet, foraging location, foraging strategy, body size and shape characteristics, and range size.

**Figure 2.**
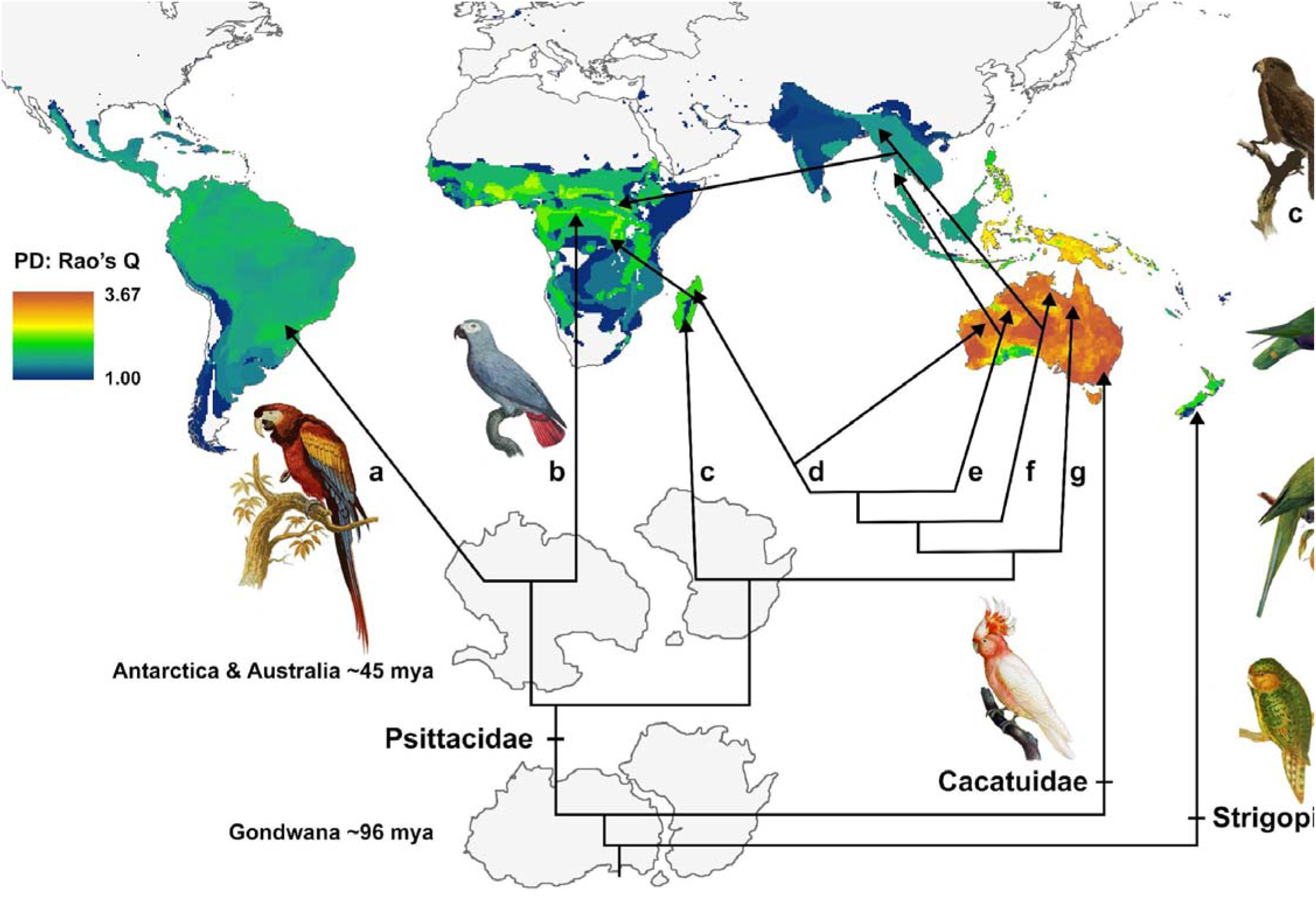
Global map of phylogenetic diversity (PD) of parrots (Rao’s Q, based on Hill numbers) associated with a diagrammatic representation of the diversification of major clades and their biogeographic affinities. Images of Gondwanaland were redrawn from Li and Powell (2001), with paths of dispersal obtained from Schweizer et al. (2010). Subfamilial designations on the cladogram are: (a) Arinae, (b) Psittacinae, (c) Coracopseinae, (d) Agapornithinae, (e) Loriinae, (f) Platycercinae, and (g) Psittaculinae. All parrot images are from the Public Domain.

IBI is highest in Australia and New Guinea (Fig. 3), and moderate in northern and central South America. For example, in South America, species richness is highest in the Amazon Basin (Fig. 4a), phylogenetic diversity is fairly evenly distributed throughout the continent (Fig. 4b), and functional diversity is highest in the dry Chaco (Fig. 4c). Although IBI equally weights each of the three dimensions (Fig. 4d), considerable spatial mismatches exist between hot spots of species richness and IBI (Figs. 4e, S1).

**Figure 3.**
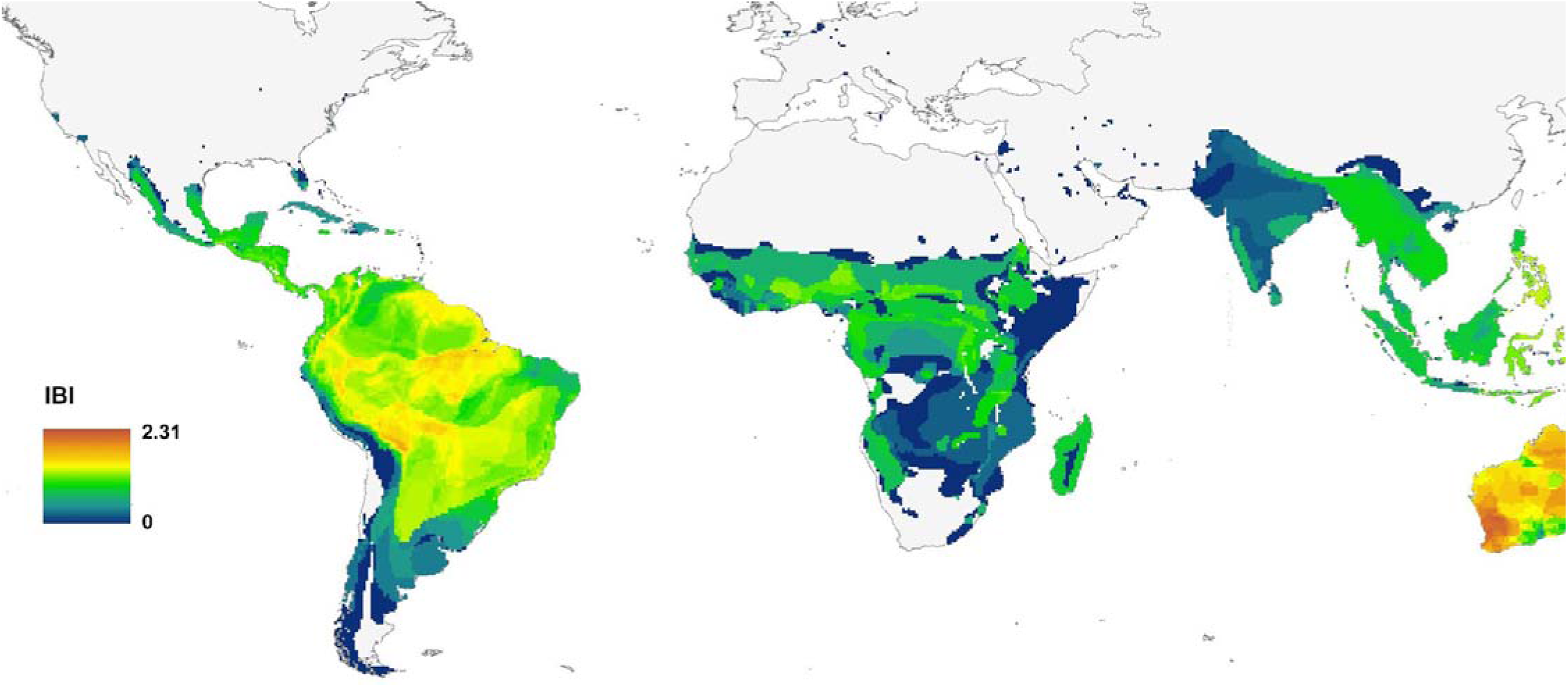
Global map of the Integrated Biodiversity Index (IBI) for parrots, which is the sum of numbers equivalents of species richness, phylogenetic diversity, and functional diversity for each grid cell, with each of the three components scaled from 0 to 1.

**Figure 4.**
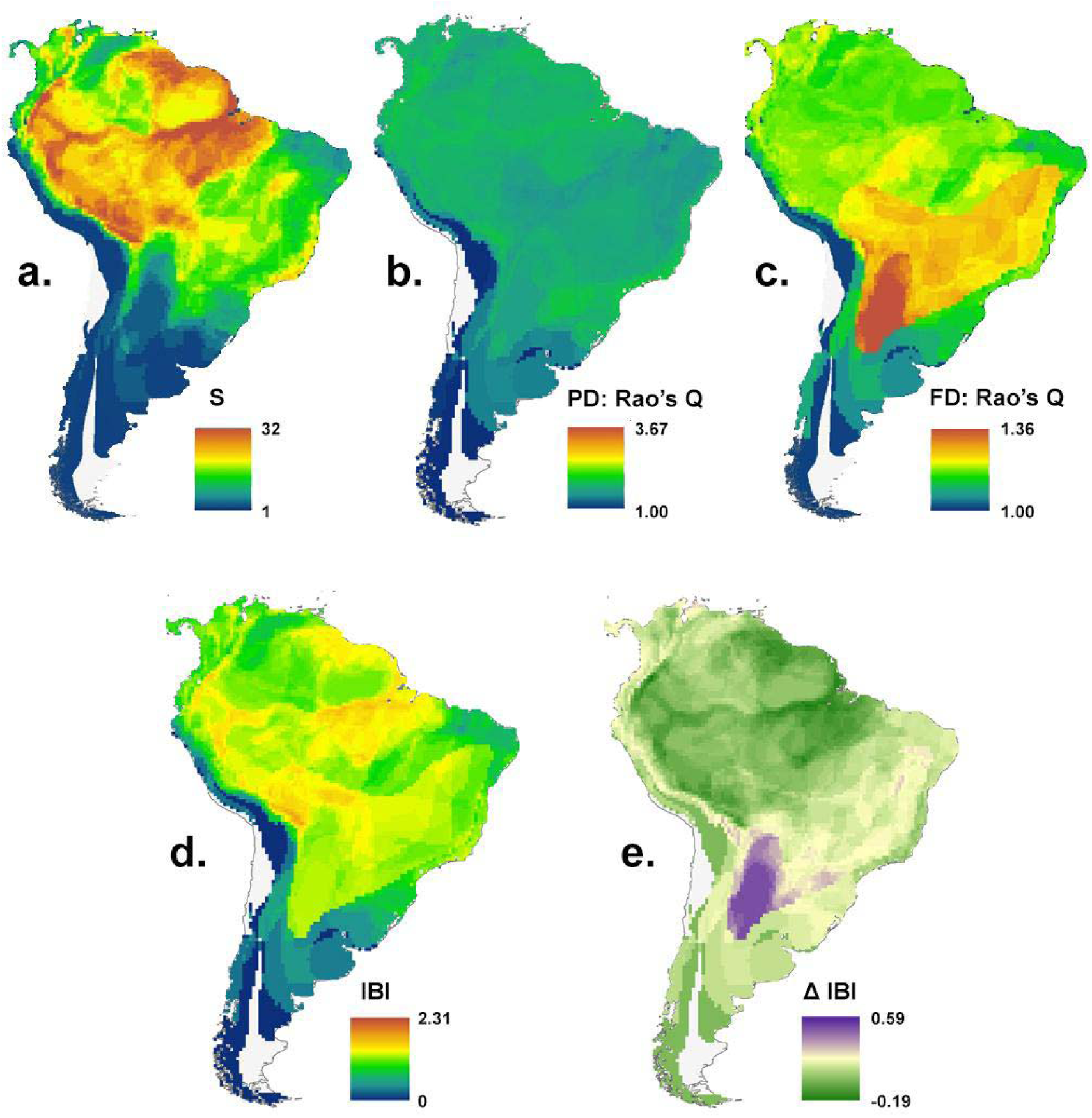
Graphical comparison of (a) species richness (S), (b) phylogenetic (PD), and (c) functional (FD) diversity patterns as well as (d) IBI of South America and (e) the difference between species richness (S) and IBI (∆ IBI), illustrating that correlation on a global level does not predict congruence of hotspots of each dimension at smaller spatial scales. To calculate ∆ IBI, we scaled the results of panels (a) and (d) to 0-1 (to make them comparable) and subtracted S from IBI, resulting in ∆ IBI, which can range from −1 to 1. Positive scores (purple) are areas more emphasized by IBI, whereas negative scores (green) are areas more emphasized by species richness. Yellow scores indicate approximately equal emphasis.

## DISCUSSION

In general, most of Australia, the island of New Guinea, and to a lesser extent, the Amazon Basin, evince the highest values of IBI. Because parrots and regions with the highest levels of species richness are generally understudied (Brito & Orpea 2009, Ducatez & Lefebvre 2014, Wilson et al. 2016), the results of IBI analysis helps to focus future research on parrots in areas of greatest need.

Aside from multiple dimensions of biodiversity, considerations of spatial scale are necessary for effective conservation planning. For example, most conservation agencies are regional or local in scale, and cannot engage in global conservation planning. For instance, the Neotropics score rather low (Fig. 2) in phylogenetic diversity compared to other regions because only one subfamily (Arinae) is endemic there. However, functional diversity is highest in the Chaco region of South America, likely because it is a harsh environment with low productivity; it likely cannot support multiple populations that perform similar functions. The Arinae subfamily diversified relatively quickly (Davies et al. 2007, Wright et al. 2008) and is the most species-rich subfamily in the parrot phylogeny, accounting for the discrepancies between dimensions of biodiversity in South America (Fig. 4). Maps of functional and phylogenetic diversity generated using only species of parrots found in the Neotropics (i.e. the Arinae) likely would identify different areas of continental conservation concern than those presented here (Figs. 2 & 4).

The areas we identified as high priorities generally correspond with results from other global prioritization research, but with a few notable exceptions. For instance, Myers et al. (2000), who initiated the “hotspots” concept, and included a wide variety of taxa, also identified Brazil’s Cerrado and the southern expanse of the tropical Andes as areas of high priority; but did not prioritize central Australia. Recent research has incorporated spatial prioritization and multiple dimensions of diversity. For instance, high priority areas for birds (Pollock et al. 2017) and mammals (Brum et al. 2017) are the Andes, equatorial Africa, Indonesia, and New Guinea, which coincide well the patterns of high IBI for parrots, though our results also emphasize southeastern Australia and the Amazon Basin, likely due to the unique diversification pattern of parrots.

The incorporation of socioeconomic data into conservation decisions can help anticipate new risks and adapt management targets accordingly (Armsworth et al. 2015). For example, high levels of urbanization correlate with an increased number of threatened parrot species, and a country’s GDP (per capita) increases the threat level (Olah et al. 2016, Butchart et al. 2015). Such an approach could identify areas within countries with increasing urbanization and increasing per capita GDP, with relatively high levels of diversity, to allow early intervention, before the effects of processes that lead to extinction (e.g. habitat loss and increased hunting) are irreversible.

If a conservation agency were to decide that hotspots of species richness were sufficient to set priorities for parrot conservation in South America, they would focus on the Amazon Basin (Fig. 4a), largely ignoring the high degree of functional diversity in the dry Chaco, which has the highest functional diversity of parrots in the world (Fig. 4c). However, by incorporating multiple aspects of biodiversity in the IBI, these aspects of biodiversity are weighted equally (Fig. 4d), allowing conservation agencies to make more informed decisions. Importantly, any particular dimension of biodiversity can be emphasized (or de-emphasized) within the IBI framework depending on the goals of a particular project. The mismatch between species richness and IBI (Figs 4e, S1) illustrates the importance of all aspects of biodiversity, and the problems with assumptions that protecting one dimension means that other dimensions are protected effectively. Spatial mismatches among hotspots of different dimensions have also been documented for mammals (Mazel et al. 2014). Conservation planners and practitioners should consider the scale and goals of conservation plans and should incorporate relevant information into an integrated framework to understand the relative value of particular policy options before taking action.

Although we do not explore extinction risk specifically, parrot species with larger ranges generally are at less risk of extinction, whereas parrot species with larger bodies or that are more dependent on forest may be at increased risk for extinction (Jones et al. 2006, Olah et al. 2016). Because these, and other traits, may be good indicators of extinction risk for parrots, it may be useful to consider each functional component separately. Mapping the areas of relatively low diversity in traits such as body size (Fig. S2), location (Fig. S4), or range size (Fig. S6), may be a good first step in identifying assemblages that may be at greater risk for extinction. Critically, simultaneous consideration of all components of functional diversity can obscure important patterns that are relevant to particular conservation issues (Spasojevic & Suding 2012, Lopez et al. 2016). Conservation agencies concerned primarily with protecting ecosystem functioning or ecosystem services may wish to focus on diet and foraging strategy diversity (Figs S3 and S5, respectively), as opposed to functional diversity.

The network of conservation areas in France provides different levels of protection for bird species richness, phylogenetic diversity, and functional diversity (Devictor et al. 2010). A potential extension of our framework could evaluate how well particular dimensions of biodiversity are protected, as a means of weighting the IBI equation to emphasize or de-emphasize particular dimensions when prioritizing areas to protect. Similarly, measures such as “ED” (Evolutionary Distinctiveness; Isaac et al. 2007), “EDGE” (Evolutionary Distinctiveness / Globally Endangered; Isaac et al. 2007), and “EDR” (Evolutionary Distinctiveness Rarity; Jetz et al. 2014) can be added or can replace phylogenetic diversity to ensure that distinct clades of the parrot tree are given more weight when assessing conservation priorities.

Climate change will have direct and indirect effects on species range shifts (Jones et al. 2016). Direct effects are based on the physiological tolerances of species, which will track their climatic niches as spatial patterns of temperature and precipitation change. Indirectly, climate change will affect land-use patterns by humans (Turner et al. 2010), which may limit or form barriers against the dispersal of individuals (Faleiro et al. 2013). Preserving connectivity among habitat patches may be a key element of effective conservation strategies in the face of climate change (Schmitz et al. 2015), further supported here by the areas we identify as high diversity, including the belt across central South America (Fig. 4). The effects of recent climate change have been greater at high elevations and in tundra compared to tropical and subtropical lowlands (Seddon et al. 2016) that harbor most species of parrots. Nonetheless, many parrot species occur in areas that are sensitive to climate change. Based on a combination of species richness and the number of threatened species and endemic species, Indonesia, Brazil, Australia, Colombia, and Bolivia are the five highest priority countries for parrot conservation action (Olah et al. 2016).

Given the predicted extent and severity of effects of climate change, conservation agencies face a daunting task. Conservation planning must balance current protection needs with future expectations as species may become locally extinct, shift their ranges, or adapt to changing conditions, possibly leading to the production of novel assemblages. Additional complications for future conservation efforts include the push and pull between different scales of conservation prioritization (i.e. the “actors” versus the “stage”; Tingley et al. 2014). Although some conservation agencies may opt to focus on particular species due to public and political values (Mace 2004), IBI is an integrative and flexible index that can incorporate multiple dimensions of biodiversity, resulting in the intuitive way to assess more than just species richness in conservation planning.

## ACKNOWLEDGMENTS

We thank R. K. Colwell, C. Rittenhouse, M. Rubega, B. Walker, and an anonymous reviewer for providing valuable feedback. KRB was supported by National Science Foundation (NSF) grant #DGE-0753455. Many of the methods in this paper were developed in part by participation of KRB, LMD, LMC, BTK, SJP, and MRW in an NSF-funded project entitled “The Dimensions of Biodiversity Distributed Graduate Seminar” awarded to S. Andelman and J. Parrish (DEB-1050680). SJP and MRW were supported by the Center for Environmental Sciences & Engineering at the University of Connecticut, as well as by an NSF award (DEB-1546686 and DEB-1831952).

