## Supplemental Information for "Integrating multiple dimensions of biodiversity to inform global parrot conservation"

**SUPPORTING INFORMATION**

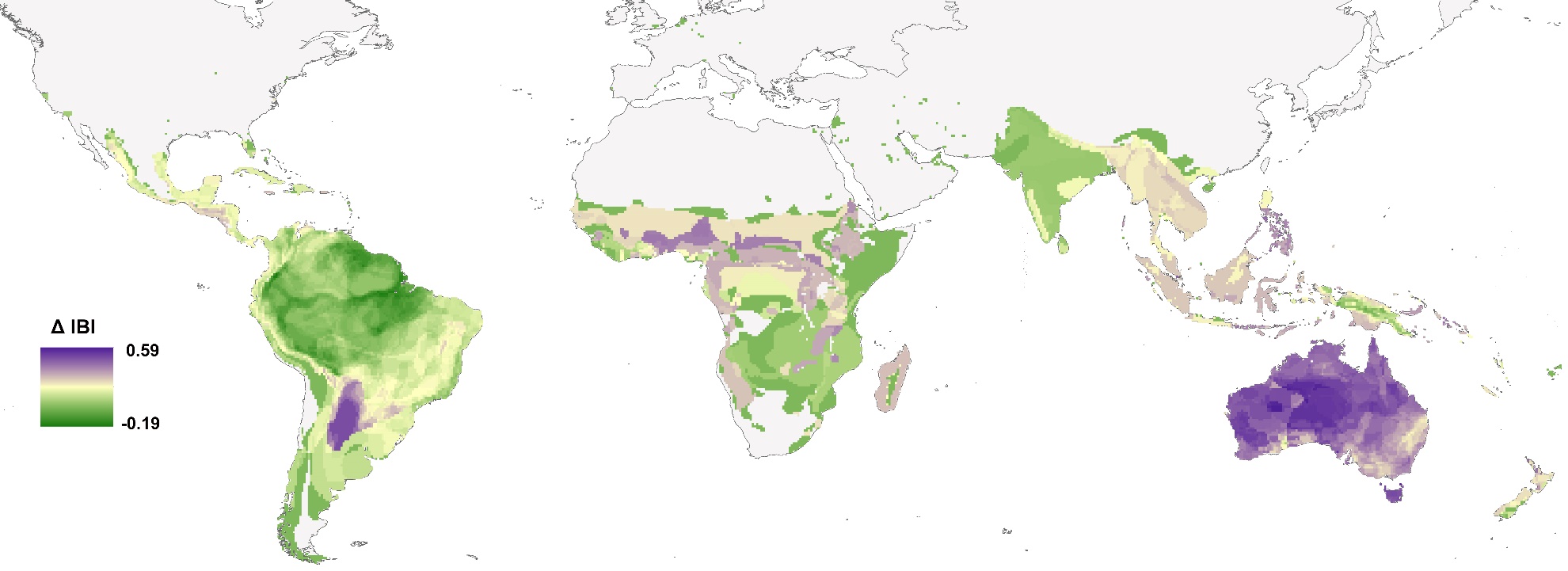

**Figure S1.** Global map of mismatch between Integrated Biodiversity Index (IBI) and species richness. Results shown in Fig. 1a and Fig. 3 are scaled from 0 to 1 (to make them comparable), and subtracted species richness (S) from (IBI), resulting in ∆ IBI*,* which can range from -1 to 1. Positive scores (purple) are areas more emphasized by IBI, whereas negative scores (green) are areas more emphasized by species richness. Yellow scores indicate similar conclusions based on IBI or species richness.

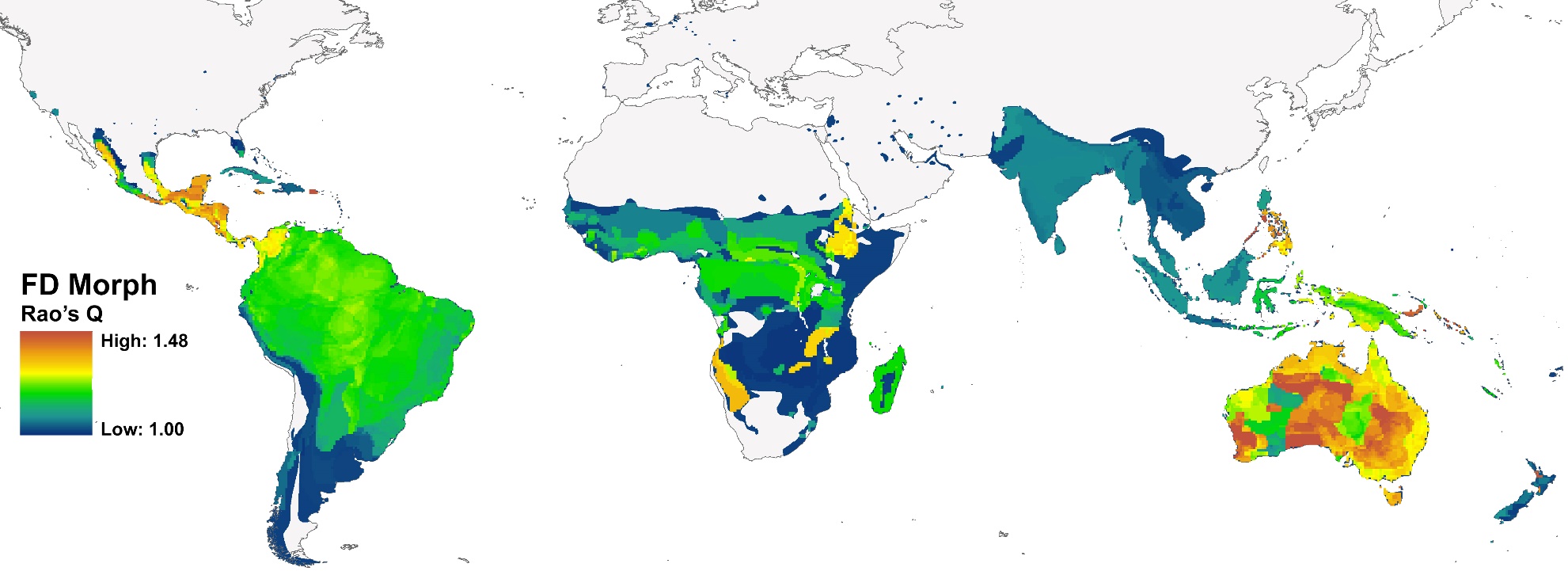

**Figure S2.** Global map of morphological trait diversity of parrots based on Rao’s Q.

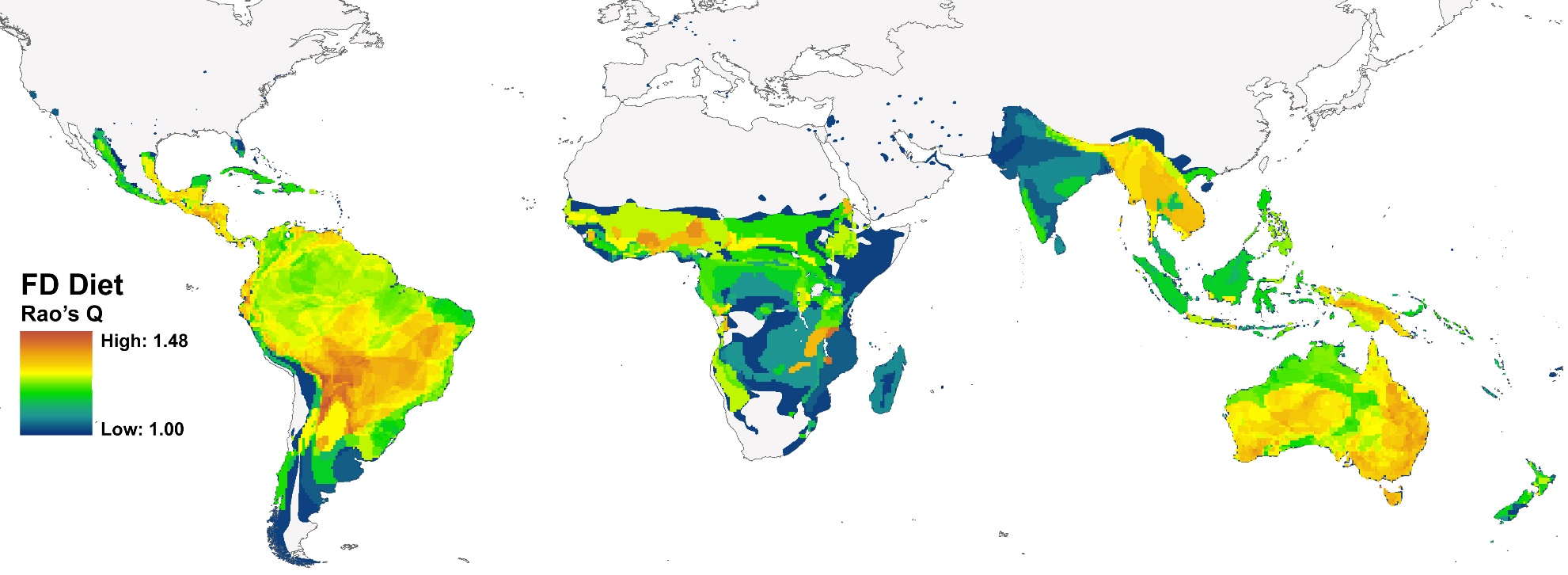

**Figure S3.** Global map of diet diversity of parrots based on Rao’s Q.

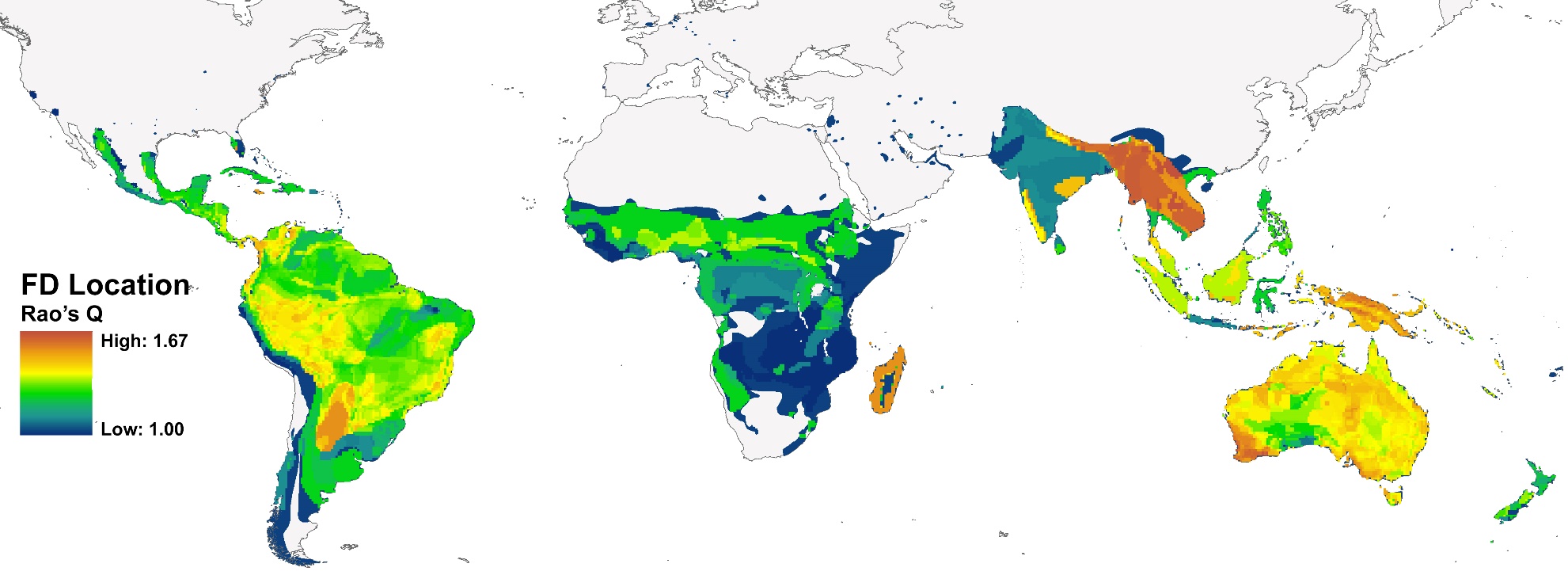

**Figure S4.** Global map of foraging location diversity of parrots based on Rao’s Q.

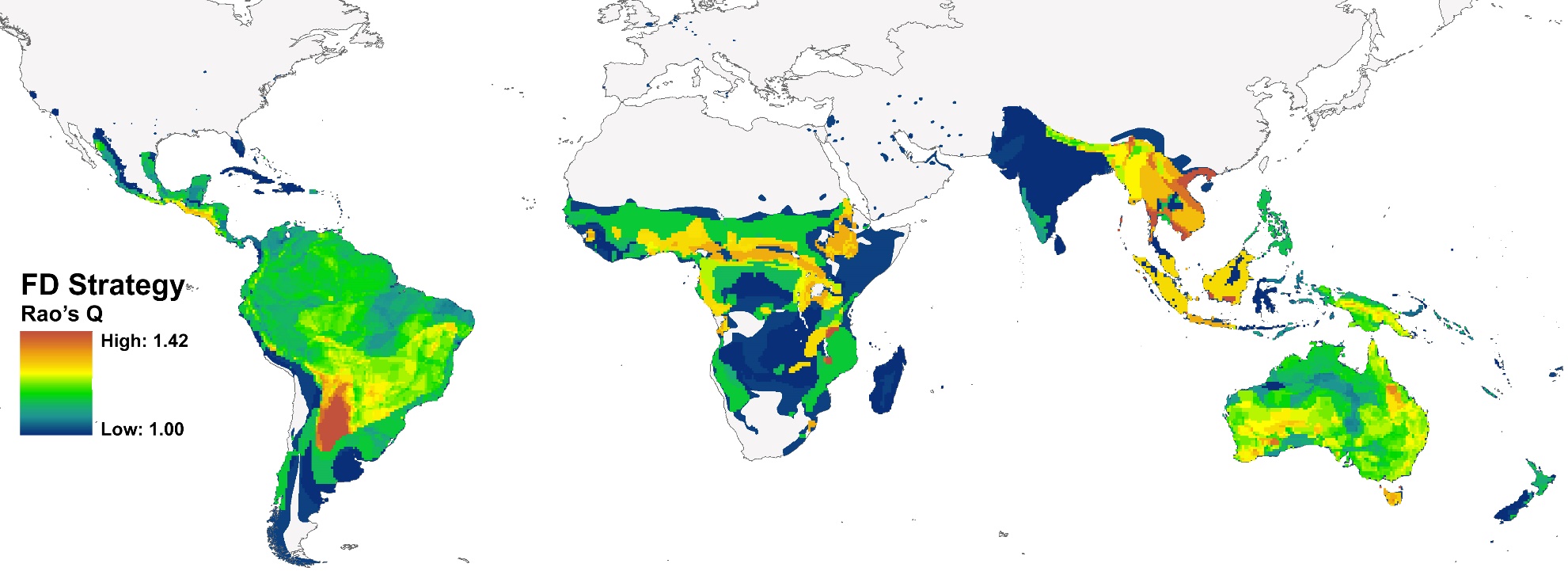

**Figure S5.** Global map of foraging strategy diversity of parrots based on Rao’s Q.

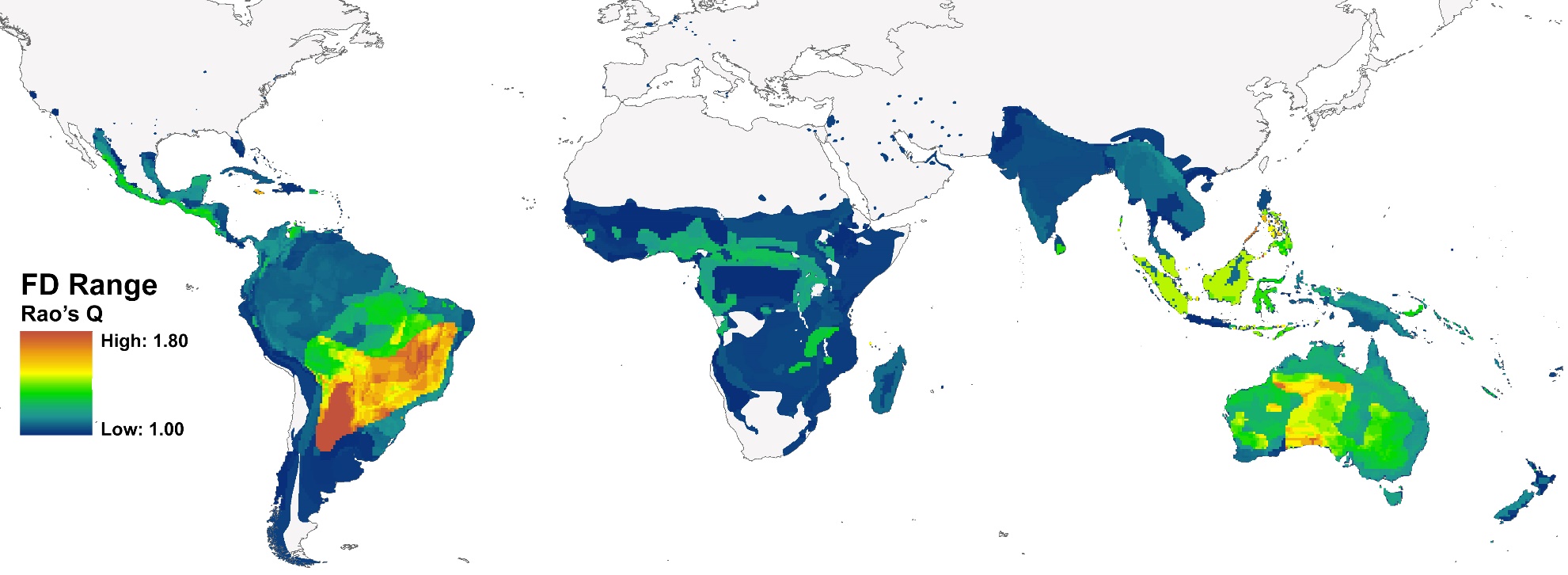

**Figure S6.** Global map of range size of parrots based on Rao’s Q.
